# Climate fluctuations drive the recruitment and growth of temperate grassland plants

**DOI:** 10.1101/2021.03.08.434383

**Authors:** Jiri Doležal, Jan Altman, Veronika Jandová, Milan Chytrý, Luisa Conti, Francisco E. Méndez-Castro, Jitka Klimešová, David Zelený, Gianluigi Ottaviani

## Abstract

Recent climate warming is associated with the increasing magnitude and frequency of extreme events, including heatwaves and drought periods worldwide. Such events can have major effects on the species composition of plant communities, hence on biodiversity and ecosystem functioning. Here we studied responses of Central European dry grassland plants to fluctuating temperature and precipitation over the last thirty years with monthly temporal resolution. We assessed the seasonal and annual dynamics of plant recruitment and growth based on the analysis of annual growth rings from the root collar. Although most studies so far applied such methods to trees and shrubs, we focused on typical grassland plants, two forbs and two chamaephytes. We related the recruitment and annual growth to monthly and annual precipitation, temperature and aridity between 1991 and 2019. We revealed species-specific responses, namely the (i) recruitment of deep-rooted, heavy-seeded species was positively affected by precipitation in both late winter-early spring and summer, whereas recruitment of shallow-rooted, light-seeded species was weakly influenced by climate fluctuations; (ii) growth of shallow-rooted species was more adversely affected by high summer temperature and drought than the growth of deep-rooted species. The population age structure of all the studied species was affected by the climate of the past decades. Most individuals established in the wet period of the 2000s, fewer in the precipitation-poorer 1990s, and the establishment was considerably reduced in the dry and warm period of the 2010s. Our findings indicate that the change towards warmer and drier climate has a profound effect even on drought-adapted ecosystems such as temperate dry grasslands. However, plant responses to various climatic extremes are species-specific, depending on their characteristics, such as life form or rooting depth. Consequently, the ongoing and anticipated climate warming will likely result in complex changes in species composition and other ecosystem properties of temperate grasslands

## 1 INTRODUCTION

Ongoing global environmental change has led to population decline in many species, especially those with restricted ranges (Staude et al., 2020). However, the main mechanisms underlying such a decline at the level of individual populations remain poorly understood (Gea-Izquierdo et al., 2021). Climate warming, prolonged drought, air pollution, and intensive land use are causing population decline either through altered biotic interactions (Gibson & Newman, 2019; Vandvik et al., 2020) or direct adverse effects on species performance and fitness (Krab et al., 2018; Babst et al., 2019). For example, water deficiency and heatwaves can directly limit the regeneration, growth, and survival of plant species adapted to wet and cold conditions (Anderson, 2016; Liancourt et al., 2020). However, little is known about the species-specific responses of plants to environmental change drivers, such as recent accelerated warming and frequent summer droughts (Brun et al., 2020; Dolezal et al., 2020). Moreover, while responses to current climate change are extensively documented in trees and shrubs (Francon et al., 2020; Altman et al., 2020), comprehensive assessments of long-term regeneration and growth responses to climate change are still rare in herbs, yet they encompass a vast majority of the plant species globally (Dee & Stambaugh, 2019; Dolezal et al., 2021).

Although regeneration and growth are crucial processes in population dynamics (Mondoni et al., 2015), range shifts (Myers-Smith & Hik, 2018), and ecosystem functioning (Büntgen et al., 2015), accurate identification of the environmental factors that control these processes remains elusive. In most global change studies related to herbs, the responses and long-term trends are derived from snapshot plant cover resurveys with long intervals between discrete sampling dates (Staude et al., 2020) without knowing what happened in-between (e.g., cyclical fluctuation, linear shift, quadratic relationship; de Bello et al., 2020). Such approaches do not necessarily lead to identifying the real trends and main mechanisms through which the changing environment limits or boosts plant performance. These can be instead identified by analysis of long-term time-series of recruitment pulses and annual biomass increments. However, such information is scarce due to technical limitations associated with the collection and analysis of well replicated and sufficiently long, annually resolved field records (Evers et al., 2021).

The retrospective anatomical determination of annual growth rings may circumvent some of these challenges (von Arx et al., 2016). Most perennial dicot plants produce annual growth rings in their belowground organs, including their oldest part, namely the root collar connecting the root and shoot systems (Klimešová et al., 2019). Such long-term radial growth series operate as “biological data-loggers” that provide this much-needed high-resolution information on recruitment, age structure and growth histories. This makes it possible to directly trace past impacts of environmental fluctuations on plant performance several decades back (Büntgen et al., 2015). Nevertheless, to our knowledge, most research so far has focused on tree species, while only a few studies explored the potential of perennial dicot herbs in capturing the effects of environmental changes and extreme events on plant recruitment and growth dynamics (Dolezal et al., 2021).

Central Europe has experienced high average temperatures and low precipitation over the last three decades (IPBES, 2019). The 1990s–2010s were the warmest interval in the last 150 years and one of the longest drought periods during the last millennia (Coumou & Rahmstorf 2012; Williams et al., 2020). It remains largely unexplored how the long-term warming trends, coupled with extreme climatic events, including frequent heatwaves and protracted summer droughts, have affected the regeneration and growth of grassland plants. Fischer et al. (2020) showed that prolonged summer droughts combined with high summer temperatures decrease grassland cover and the dominance of polycarpic perennials while increasing the abundance of short-lived ruderal species. However, it remains unclear which major mechanisms at the level of plant populations and individuals are behind these climate-induced and widespread responses. Additional studies are needed to clarify whether the climate-induced population declines mainly result from reduced germination and seedling survival (Lu et al., 2019) or limited growth, fecundity and increased mortality of adult plants (Brun et al., 2020). In Central European temperate grasslands, no study had simultaneously followed the long-term dynamics of recruitment and radial growth of individual species in natural conditions to determine the climatic factors associated with demographic trends and growth fluctuations. The current study is the first attempt to bridge this knowledge gap for climate change science and functional ecology in the region.

We selected four temperate dry-grassland plant species with relatively long lifespan (up to 30 years) and distinct annual rings, making age dating possible (Schweingruber et al., 2011). We collected demographic and annual growth data for populations of two perennial forbs (*Lychnis viscaria, Silene nutans*) and two chamaephytes (*Helianthemum grandiflorum, Thymus pulegioides*), which are specialists of rock-outcrop grasslands. Such grasslands constitute an extreme habitat in Central Europe because of high irradiance coupled with shallow and relatively nutrient-poor soils. Therefore, we expected the recruitment, age structure and growth of the focal specialists of this extreme habitat to be sensitive to environmental fluctuation. We also expected species-specific reactions to climate change drivers due to differences in growth forms (Klimešova, 2018), xylem anatomy (von Arx et al., 2016), rooting depth (Kutschera & Lichtenegger, 1992), and seed mass (Kleyer et al., 2008).

Specifically, we ask: (Q1) Which climatic factors (temperature, precipitation, drought index) correlate with recruitment and growth dynamics? (Q2) How much is the radial growth of the studied species sensitive to climatic fluctuations, recent warming trends, and extreme climate events? (Q3) Do high temperatures limit recruitment and growth directly, or rather via drought caused by increased evapotranspiration under warmer conditions? (Q4) Do species with contrasting life histories (forbs versus chamaephytes, deep-rooted versus shallow-rooted plants) show specific responses to weather fluctuations? (Q5) Do current age distributions suggest population contraction or expansion? The fundamental expectation is that wet springs and cool summers tend to increase recruitment because seed germination and seedling survival are primarily constrained by dry springs and summer heatwaves in Central European dry grasslands. We also hypothesized that radial growth is promoted by late spring and summer precipitation while it is reduced by high summer temperatures. We also expected the recruitment to be more contingent on precipitation in species with heavier than lighter seeds, and the growth to be more sensitive to summer heatwaves in shallow-rooted than deep-rooted species.

## 2 METHODS

### 2.1 Study area

The study area is located in the surroundings of Třebíč, western Moravia, Czech Republic (geographical coordinates N 49.21–49.27°, E 15.91–16.02°; elevation 430–480 m a.s.l.). It experiences a temperate climate (mean annual temperature = 6.5–8 °C, total annual precipitation = 500–550 mm; Tolasz, 2007) characterized by a remarkable seasonality, namely cold winters with January means below 0 °C and warm summers. Precipitation maxima occur in summer, but there can be prolonged drought spells lasting for a few weeks in spring or summer. Based on previous studies (e.g. Zelený & Li 2008), we selected 20 dry grassland sites, occurring as terrestrial habitat islands surrounded by arable land, across an area of approximately 100 km^2^. These grassland patches develop on low dome-shaped elevations on granitic outcrops (granite or syenite). The size of these grassland patches ranges between 361 m^2^ and 14,115 m^2^ (mean = 2,912 m^2^), and their elevations are usually less than 10 m above the surrounding landscape. Several soil parameters were collected at the grassland level, i.e. 2 to 3 randomly selected points at each site (depending on site area), in the same location where the automatic TMS3 microclimatic stations were positioned (see below). The mean soil depth across sites is approximately 9.5 cm (average obtained from > 2500 soil depth measurements across the 20 sites). The soils are mostly derived from the weathering of the granitic bedrock, hence characterized by a coarse-grained texture (mean sand content = 81%), low water retention capacity and pH (mean = 4.7), low nutrient contents (mean total nitrogen = 0.76%; mean plant-available phosphorus = 90.5 mg/kg), and soil organic carbon (mean = 9.4 mg/kg). The grasslands are former low-productive pastures, now abandoned for several decades, and some of them managed for conservation purposes, mostly through mowing They belong to phytosociological alliances *Hyperico perforati-Scleranthion perennis* (class *Koelerio-Corynephoretea*) and *Koelerio-Phleion phleoidis* (class *Festuco-Brometea*) (Chytrý, 2007; Zelený & Li, 2008).

### 2.2 Focal species and data collection

During summer 2019, we collected 174 individual plants belonging to four focal grassland species adapted to the dry conditions of shallow and sandy soils on the granitic outcrops. At each site, we collected three randomly selected individuals per species. These species included the forbs *Lychnis viscaria* (Caryophyllaceae; 57 individuals) and *Silene nutans* (Caryophyllaceae; 42), and the chamaephytes *Helianthemum grandiflorum* subsp. *obscurum* (Cistaceae; 18) and *Thymus pulegioides* (Lamiaceae; 57) (Figure 1). The lower number (i.e. less than 60 individuals/species) was because not all the species occurred across all the study sites.

**Figure 1.**
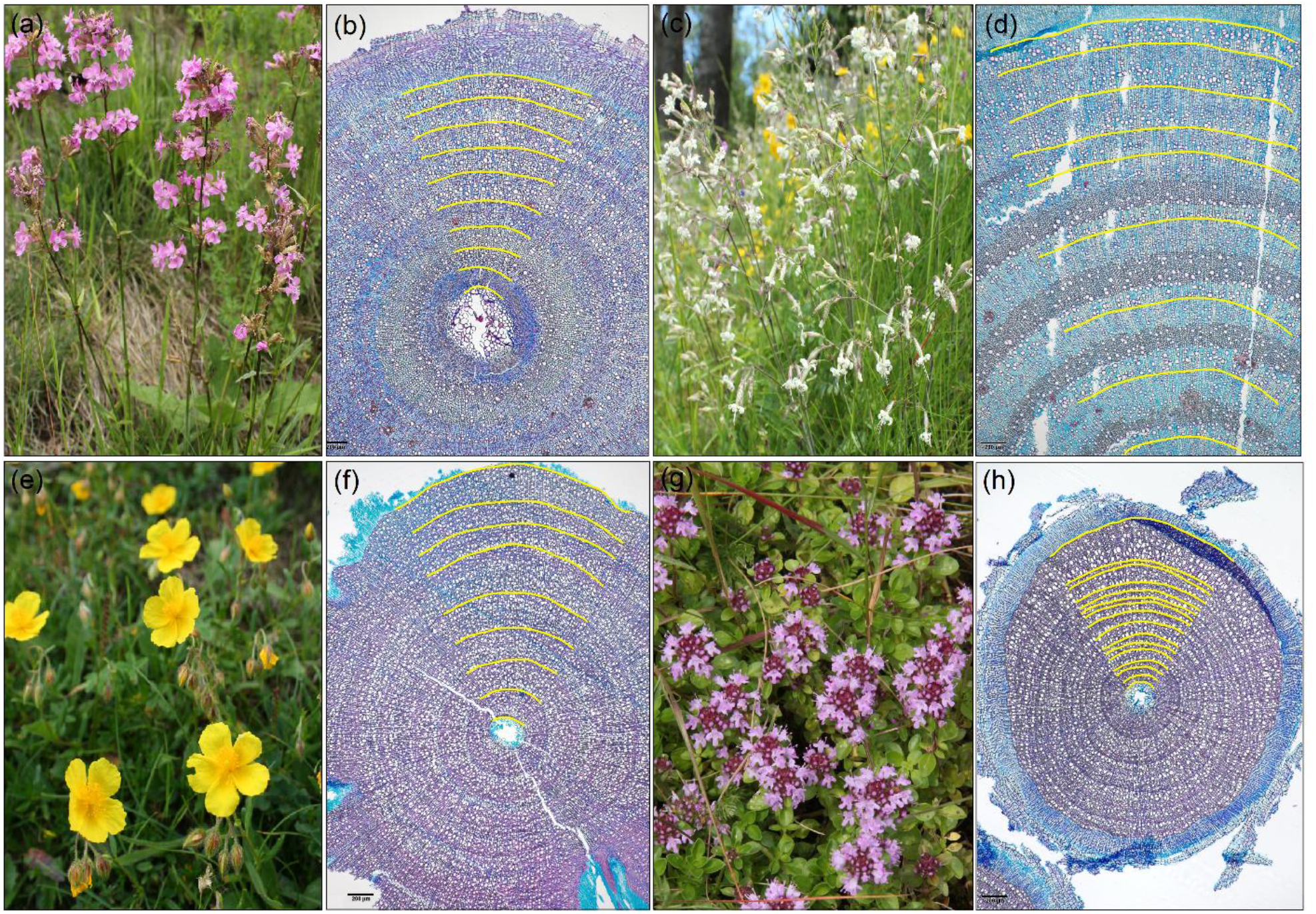
Studied species and their double-stained cross-sections of root collars with growth- rings (highlighted by yellow lines): (a, b) *Lychnis viscaria*, (c, d) *Silene nutans*, (e, f) *Helianthemum grandiflorum* and (g, h) *Thymus pulegioides*.

These four species are suitable for age dating and growth analysis because of distinct annual rings characterized by ring-porous (*Thymus, Lychnis*) or semi-ring porous (*Silene, Helianthemum*) xylem structures with marked differences between earlywood (40-60 µm in diameter) and latewood (5-10 µm) vessels (Figure 1), thick-walled and/or radially flattened latewood fibers (*Helianthemum, Thymus*), distinct latewood fiber zone (*Helianthemum*), and a continuous tangential band of latewood parenchyma (*Lychnis, Silene*) (Table 1). Absolute dating is possible due to the rarely rotten stem (unlike in other grassland plants), while cross-dating (assigning the calendar years to individual annual increments) is facilitated by the general lack of missing rings.

**Table 1.** Characteristics of the studied species.

|  | *Lychnis viscaria* | *Silene nutans* | *Helianthemum grandiflorum* | *Thymus pulegioides* |
| --- | --- | --- | --- | --- |
| Number of individuals | 57 | 42 | 18 | 57 |
| Individuals used in chronology | 21 | 17 | 8 | 19 |
| Inter-series correlation | 0.44 | 0.29 | 0.43 | 0.40 |
| Chronology agreement (GLK in %) | 60 | 61 | 67 | 56 |
| Mean and max. plant age (yr) | 11-22 | 14-29 | 15-25 | 17-29 |
| Plant height in maturity (cm) | 30-75 | 30-60 | 10-40 | 10-30 |
| Rooting depth (cm)<sup>1</sup> | 90 | 160 | 160 | 60 |
| Seed mass (mg)<sup>2</sup> | 0.0744 | 0.298 | 1.381 | 0.1964 |
| Flowering period<sup>3</sup> | May-Jun | May-Jul | Jun-Sep | Jul-Oct |
| Mean radial growth (µm) | 367 | 350 | 300 | 393 |
| Max. vessel size (µm<sup>2</sup>) | 711.2 | 663.7 | 449.5 | 632.1 |
| Min. vessel size (µm<sup>2</sup>) | 110.0 | 124.5 | 164.0 | 178.4 |
| Cross-sectional vessel area (%) | 23% | 22% | 21% | 24% |
| Vessel number per 1 mm<sup>2</sup> | 743 | 712 | 711 | 686 |
| Lignified tissue (%) | 21.1 | 15.4 | 46.1 | 38.4 |
| Parenchymatic tissue (%) | 51.9 | 58.8 | 30.5 | 31.0 |
| Conductive tissue (%) | 27.0 | 25.8 | 23.4 | 30.6 |

1 – Kutschera & Lichtebnegger, 1992; 2,3 – Kleyer et al., 2008.

To obtain age and growth data for each individual, a segment of the oldest root portion (about 5 cm long) was excised from each plant and placed in 40% ethanol to keep the root tissue soft and prevent mold development. In the laboratory, we followed standardized procedures (Gärtner & Schweingruber, 2013). We cut several cross-sections for each individual from the oldest plant tissues between the hypocotyl and the primary root (root collar) using a sledge microtome, stained with Astra Blue and Safranin and permanently fixed on a microscope slide with Canada Balsam (Dolezal et al., 2018). All annual rings of perennial dicot plants should exist in the root collar zone (Büntgen et al., 2015). High-resolution images of these microsections were taken using an Olympus BX53 microscope, Olympus DP73 camera and the cellSense Entry 1.9 software. We then measured the annual radial growth increments. The age of each individual was estimated as the maximum number of annual rings counted along the four radii for each cross-section. We also recorded vessel lumen area, the number and area of vessels per unit area and the proportion of three basic tissues: lignified (mechanical support), parenchymatic (carbohydrate storage), and conductive (vessel conduits transporting water) (Table 1) (Klimešova et al., 2019).

### 2.3 Climate data

Recruitment and growth time-series were correlated with the meteorological data from the nearby weather station in Velké Meziříčí (E 16.0086; N 49.3528; Figure 2), located c. 10 km from the study sites at a similar elevation. The mean daily temperatures from this station were correlated (r = 0.97) with in-situ records of air and soil temperature that we measured at 15-minute intervals using a network of automatic TMS3 microclimatic stations (TOMST, Czech Republic; Supplementary Information Figure S1), having 2 to 3 of them positioned at each site. Therefore, the long-term meteorological data represent a reliable source of information to test the correspondence between inter-annual changes in climate and plant recruitment and radial growth of our focal species. Additionally, to investigate the impact of moist and dry years on plant recruitment and growth, we used the monthly 0.5°×0.5°-gridded Standardized Precipitation-Evapotranspiration Index (SPEI) as a proxy of wetness (positive value) and dryness (negative values), for the grid cell 49.0-49.5° N and 15.5-16.0° E (www.climexp.knmi.nl).

**Figure 2.**
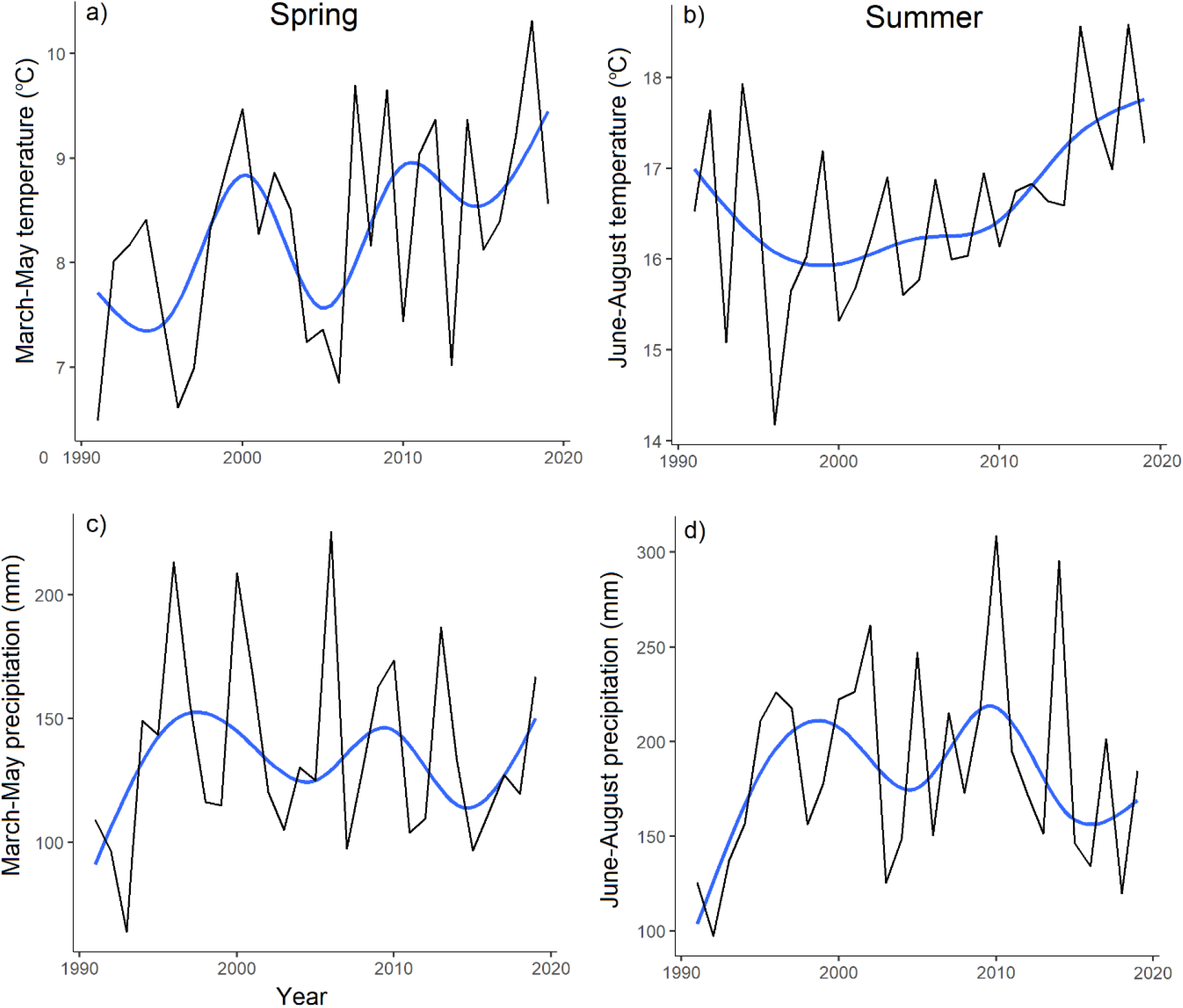
Time series of the mean (a) spring and (b) summer temperatures and (c) spring and (d) summer precipitation totals from 1991 to 2019 based on the records from a local climate station. A 6-years smoother line was fitted using the RLM (robust fitting of linear models, blue lines).

### 2.4 Data analysis: Recruitment-climate relationship

All measured series were used to assess the climatic influence on age distributions. We defined the year of plant establishment as the calendar year of the first ring (measured from the center) from the oldest series of each individual. We aggregated the number of established individuals to three-year intervals to minimize the possible impact of missing/false rings for the series which have not been cross-dated (Büntgen, 2019; Altman 2020). Precipitation and temperature data were averaged over three-year intervals. Consequently, Pearson’s correlation coefficients were calculated between establishment chronology and temperature, precipitation and SPEI. Correlations were calculated for the same period as the growth-climate relationships described below (February to September).

### 2.5 Data analysis: Growth-climate relationship

Individual annual-increment width series were first visually cross-dated and then statistically verified based on the percentage of parallel variation (GLK-Gleichläufigkeit; Table 1), which is the sign test commonly used in dendrochronology (Buras & Wilmking, 2015). The GLK values of over 60% are generally considered sufficient for the reconstruction of past climate variation (Esper & Gärtner, 2001). To remove non-climatic age-related growth trends from the individual time series, residual chronologies of individual annual-increment width series were developed using the R packages *dplR* (Bunn, 2008). A negative exponential curve or a linear model with negative or zero slope (to preserve positive trends presumably due to climate in the raw data series) was fitted to each measured series. Consequently, we built the species-specific mean growth chronology by calculating a robust mean of detrended ring width series (Bunn, 2008). The relationships between the mean growth chronologies and temperature, precipitation and SPEI were assessed based on bootstrapped Pearson’s correlation estimates. Bootstrapped confidence intervals were used to estimate the significance (p < 0.05) of the correlation coefficients (Zang & Biondi, 2015). Correlation coefficients were calculated for monthly climate variables starting in February through September (8 months).

## 3 RESULTS

### 3.1 Climate-dependency of plant recruitment

The distribution of plant ages was symmetric, Gaussian-like in *Lychnis* and *Silene*, and more skewed towards older individuals in *Helianthemum* and *Thymus* (Figure 3). In all the four species, most individuals were 10-20 years old. We found significant correlations between climate and plant establishment in three of the four species, with recruitment peaks associated with both late winter and mid-summer temperature, precipitation and SPEI. The recruitment during the past 30 years was associated with different climatic signals in different species (Figure 3), except for *Lychnis*, in which it was not significantly correlated with any of the climate variables tested. The recruitment of *Silene* was positively and significantly correlated with February and August precipitation and SPEI. Likewise, in *Helianthemum*, higher February, March and August precipitation and SPEI over the past 30 years coincided with intensified plant recruitment, while August heatwaves harmed recruitment. Therefore, the period of rainy summers in 2004-2009 triggered peak recruitment for both *Silene* and *Helianthemum*, whereas the drier and warmer summers in 1991-1995 and 2012-2019 coincided with reduced plant recruitment (Figure 3). The recruitment pulses in *Thymus* were negatively correlated with high March temperatures, with the establishment being the highest during colder springs of 2002-2009 (Figure 2) and the lowest between 2012-2019 when March temperatures were the highest in the past thirty years.

**Figure 3.**
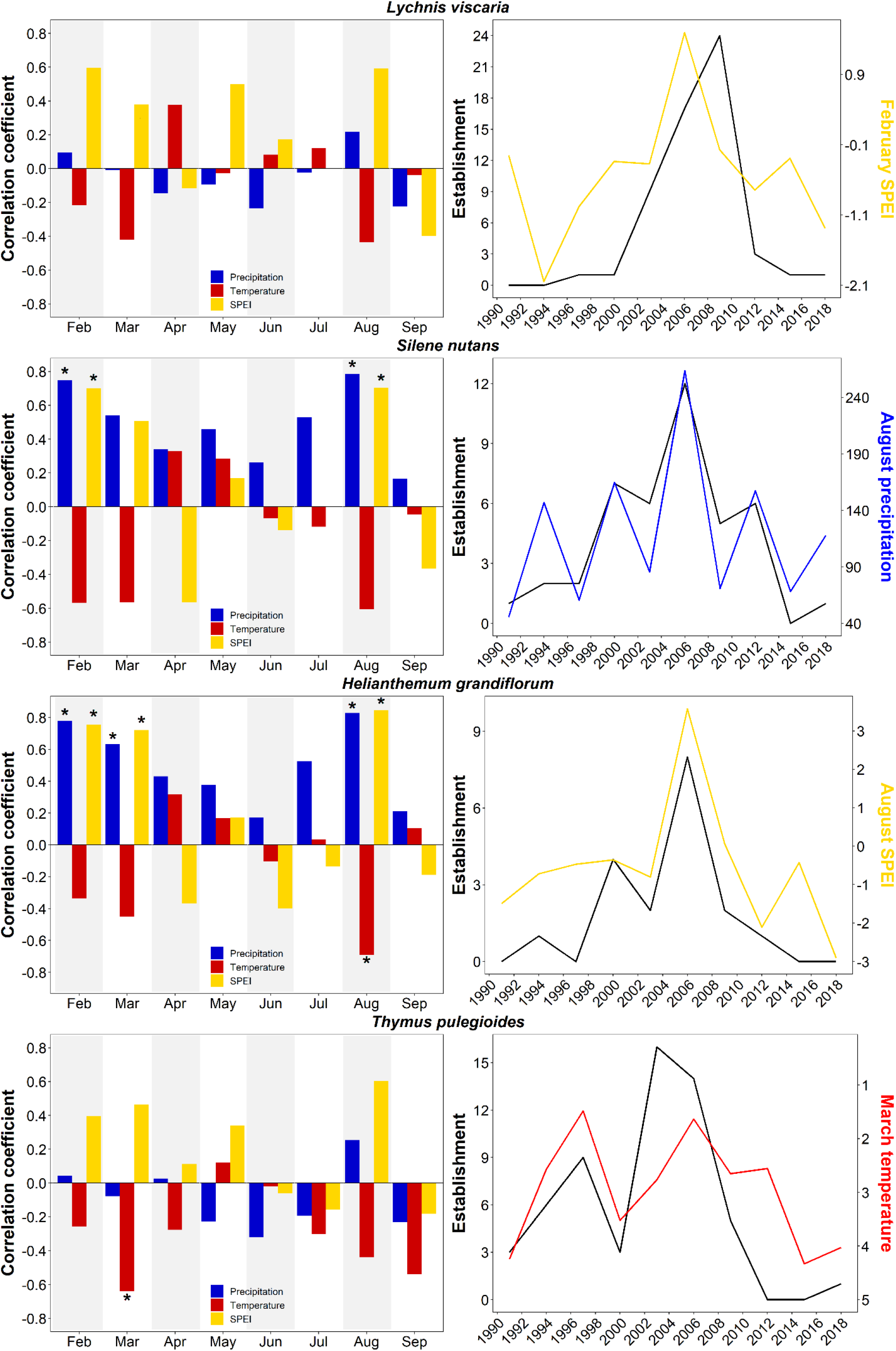
Relationship between the number (three-year sums) of newly established individuals of the focal species and monthly temperature, precipitation and SPEI index (left column; three-year averages); significant correlations are indicated by an asterisk. The right column shows the annual numbers of established individuals of the focal species (black line) compared with the climatic parameter with the strongest establishment-climate relationship (color line). Note that the temperature axis in *Thymus* is in the reverse direction.

### 3.2 Climate-dependency of plant growth

We identified contrasting climatic plant growth controls in the four focal plant species (Figure 4). *Lychnis* growth was significantly and positively correlated with July precipitation and negatively correlated with July temperatures, August precipitation and August SPEI. The wet July in 2008-2009 and 2016-2017, therefore, promoted plant growth in *Lychnis*, while hot July in 2006 and 2015 impeded plant growth. Cold and rainy late summer (August) had also a negative effect on *Lychnis* growth as in years 2002, 2006, and 2010. *Silene* growth was positively correlated with late winter (February) precipitation and temperatures and early spring (March) precipitation and SPEI. Hence, the wet March in 2006-2009 boosted plant growth in *Silene*, while dry spring in 2001-2005, 2010-2012, and 2018 reduced radial increment. *Helianthemum* growth was negatively correlated with February and July temperatures. The enhanced radial growth in *Helianthemum* coincided with low February temperatures, as in 2003, 2005, and 2012, while the reduced growth occurred in 2002, 2007, 2015, and 2016 when February temperatures were higher than average. *Thymus* growth was positively correlated with June precipitation and SPEI, and negatively correlated with June temperatures (Figure 4). Therefore, high rainfall and low temperatures in June coincided over the past thirty years with increased plant growth in *Thymus*, as in 1995, 1998, 2009-2010, 2013 and 2018, while low rainfall, high temperatures and droughts in 1994, 1996-1997, 2003, 2008 and 2011 corresponded with reduced plant growth.

**Figure 4.**
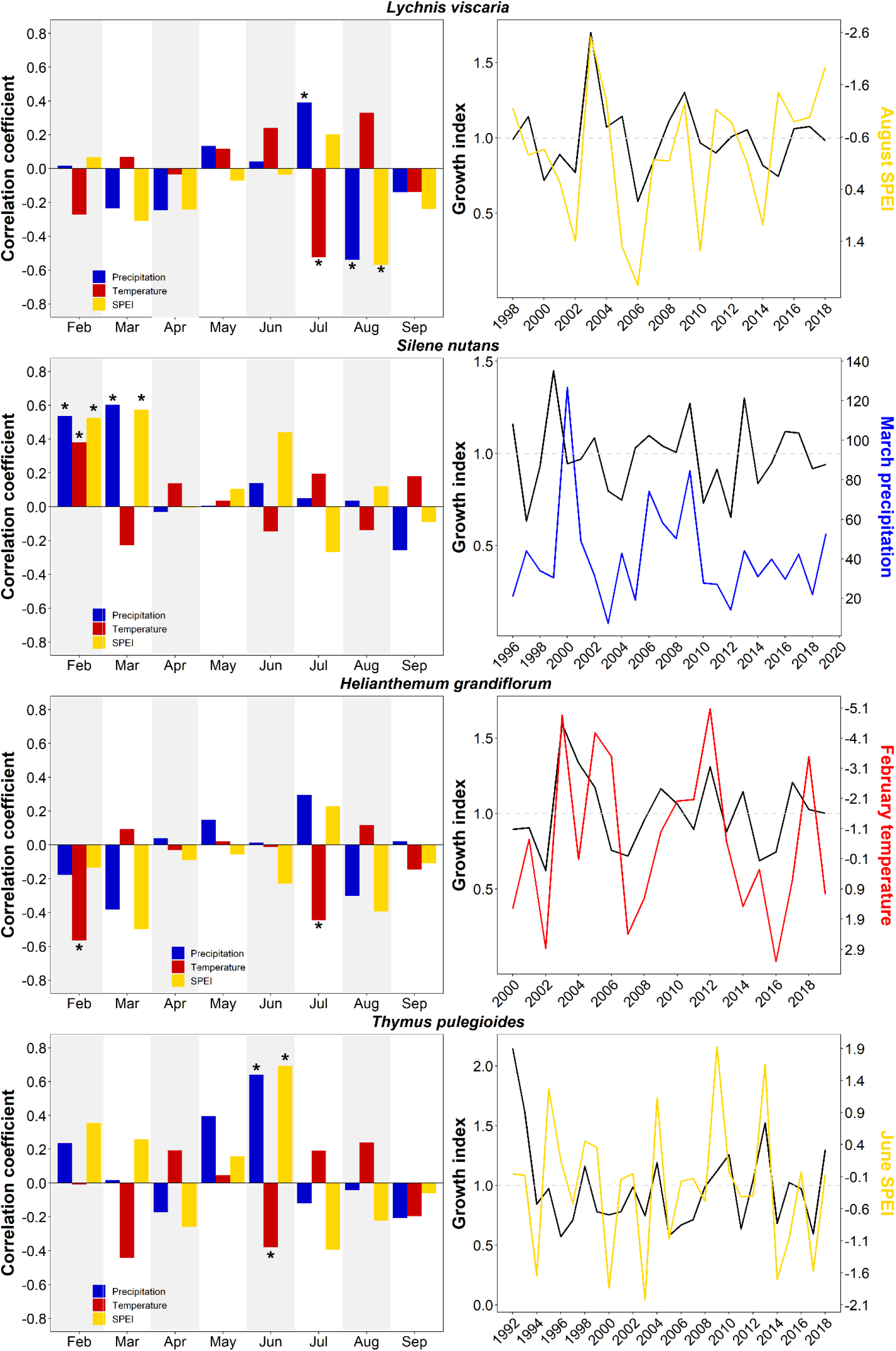
Relationship between growth indices of the focal species and monthly temperature, precipitation, and SPEI (left column); significant correlations are flagged by an asterisk. The right column compares the annual changes in the growth index and the climatic parameter with the strongest growth-climate relationship. Note that the SPEI axis in *Lychnis* and the temperature axis in *Helianthemum* are in the reverse direction.

## 4 DISCUSSION

This study constitutes one of the first attempts to examine long-term recruitment and growth dynamics in temperate grassland plants in relation to changing climate. Our results support the growing evidence showing marked sensitivity of individual plant species to global climate change drivers (Büntgen et al., 2015; Dee et al., 2018; Myers-Smith & Hik, 2018, Dolezal et al., 2020). The application of a retrospective analysis of annual growth rings in the populations of two forbs (*Lychnis viscaria, Silene nutans*) and two chamaephytes (*Helianthemum grandiflorum, Thymus pulegioides*) from 20 temperate dry grassland sites in Central Europe provided novel insights into the mechanisms underlying the long-term recruitment and growth processes in rapidly warming and drying European temperate grasslands. By sampling populations across different sites and species, we identified contrasting climatic controls during recruitment and growth for different species, as well as demographic trends emerging from age distributions. These findings stress the complexity of plant responses to climate change and reinforce the need to focus on species-specific responses when predicting climate change impacts on populations (Rixen et al., 2010; Bellard et al., 2012). Such insights have multiple implications for understanding ecosystem functioning and diversity (Evers et al., 2021).

### 4.1 Recruitment and population dynamics under climate change

This study clearly shows that recent climate warming and prolonged drought events exerted a strong negative impact on grassland plant recruitment, growth and population dynamics, which is consistent with previous observations from trees (Altman et al., 2020; Babst et al., 2019), shrubs (Krab et al., 2018; Buchwal et al., 2020) and alpine herbs (Dee & Stambaugh 2019; Dolezal et al., 2020, 2021). In Central Europe, the last three decades can be divided into warmer and drier 1990s and 2010s and relatively cooler and wetter 2000s. The most favorable conditions for the establishment of new individuals occurred in the 2000s, while recruitment in the 1990s and the 2010s was significantly reduced. As a result, the populations of the four focal species have age distributions with greater numbers of individuals aged 10-20 and fewer young individuals due to reduced recruitment during the warmer and drier last decade, suggesting an aging population. This aligns with recent evidence of senescing populations from other grassland systems exposed to accelerated warming and diminishing precipitation (Zhang et al., 2020; Ronk et al., 2020; Dolezal et al., 2021).

Recruitment was especially low during extreme heatwaves as in the summers of 1992, 1994, 1995, 2003, 2015 and 2018, which had severe impacts on vegetation productivity (Ciais et al., 2005), species composition (Fischer et al., 2020) and forest dieback in Europe (Brun et al., 2020). Summertime European heat and drought waves are induced by wintertime rainfall deficit that spreads from the Mediterranean region northward throughout continental Europe due to atmospheric transport of anomalously warm and dry air (Vautard et al., 2007; Schär et al., 2004). This can also be the case of the studied central European grasslands, as confirmed by the negative correlation between February-April precipitation and August temperatures (Pearson r = -0.48). Higher frequency of hot summers preceded by winter precipitation deficits during the past 30 years may therefore explain a strong positive correlation between the plant recruitment and February-March and August precipitation and SPEI, and a negative correlation between the recruitment and August temperatures, presumably through direct impacts on seed germination during dry spring and seedling survival in hot summer (Harrison & La Forgia, 2019; Yi et al., 2019). Although the sensitivity of emergence and survival of seedlings to temperature has been well documented (e.g., James et al., 2019; Yi et al., 2019), our results emphasize the role of synergies between winter and spring drought and extremely hot summers. Such synergies may be critical for the water reservoir in the soil (Schär et al., 2004), with implications for plant regeneration and the maintenance of stable populations (Evers et al., 2021).

However, the positive effect of both late winter-early spring and summer precipitation on recruitment was not common to all focal species, as reported in other studies (e.g., Walder & Erschbamer, 2015). This was particularly evident in deep-rooted, heavy-seeded species (*Helianthemum* and *Silene*), whereas recruitment of shallow-rooted, light-seeded species (*Lychnis* and *Thymus*) was weakly affected by precipitation fluctuations. Drought and heat are critical for seed germination and juvenile growth (Reader et al., 1993), so plants with deeper roots and heavier seeds may cope better with climate extremes by reaching possibly stable water table sources in the deeper soil layers (Schenk & Jackson, 2002), and through larger carbohydrate and nutrient storage in seeds to promote seedling growth (Mašková & Herben, 2018).

### 4.2 Species-specific climate controls of plant growth dynamics

Unlike the differences in recruitment dynamics, which were mainly driven by fluctuations in late winter and late summer precipitation, plant growth depended on early summer temperatures and rainfalls. We found significant declines in growth with increased temperature in June-July and reduced precipitation between June and August, similarly as in the Rocky Mountain (Dee & Stambaugh, 2019) and Himalayan (Dolezal et al., 2021) alpine plants. Similarly to recruitment, we found inter-annual growth dynamics to be dependent on multiple climatic variables, indicating the crucial role of synergies between summer drought and heat. Plants exposed to drier air transpire more water, emptying the soil moisture reservoir (Vautard et al., 2007). Drier soils then emit more sensible heat, inhibiting cloudiness and further increasing daytime shortwave radiation and temperature (Schär et al., 2004). Dry soils further increase evapotranspiration, which induces stomatal closure to prevent cavitation and desiccation of leaf tissues. Harsh conditions then restrict cellular processes, including cell division, cell enlargement, and cell differentiation (see Figure 1), leading to smaller and fewer earlywood vessel conduits (Schweingruber et al., 2011), which diminishes plant capacity for water transport and impairs carbon fixation (Sperry, 2003). This is finally reflected in narrow growth rings and annual biomass production (Dolezal et al., 2018).

We identified species-specific climatic controls of inter-annual growth fluctuations presumably due to specific ecological (life history) strategies linked to contrasting species morphological and anatomical structures and functions. For example, the growth of shallow-rooted *Thymus* was promoted by high May-June precipitation and reduced by high June temperatures. May-June precipitation was the most important growth driver in *Thymus*, similarly to other species with semi-ring porous xylem, typically growing in habitats with wet spring and dry summers such as in the Mediterranean biome. In such conditions, where the early growing season is relatively cold and moist while the late-season is warm and dry, plants tend to develop variable vessel conduit sizes in growth rings (Schweingruber et al., 2011), which change characteristically from large earlywood to small latewood cells, reflecting the dominant impact of either low temperature or water shortage on the seasonal growth-ring formation (Dolezal et al., 2019). By shifting the onset of growth activity and large earlywood vessel formation from early spring to May-June, *Thymus* may avoid spring frost damages to fully rehydrated tissues, while the formation of smaller vessels later in the growing season may prevent drought-induced cavitation. Semi-ring porous xylem with wide earlywood vessels in spring and narrow latewood vessels in summer, therefore, should provide both efficiency and safety of the water transport as well as a potentially longer growing season (Schweingruber et al., 2011; von Arx et al., 2016; van der Sande et al., 2019).

*Lychnis* growth was dependent on climatic conditions later in the growing season, stimulated by high July precipitation and harmed by high July temperatures and August precipitation. This may have affected the formation of latewood bands of marginal parenchyma, hence plant storage ability and next-spring regrowth capacity (Schweingruber et al., 2011). Favorable July and August conditions may promote additional cell division leading to the wider latewood parenchyma band and hence larger total annual growth increment. Unlike the studied chamaephytes, which have lignified rootstock and perennial woody branches, the studied forbs must recover annually most of the aboveground biomass lost during winter dormancy. The spring regrowth relies heavily on sugar stored in latewood parenchyma cells of belowground organs (Plavcová & Jansen, 2015). This is reflected in a twice as high proportion of storage parenchyma tissue in the studied forbs compared to chamaephytes (Table 1). Indeed, in our chamaephytes, *Thymus* and *Helianthemum*, thick-walled fibers (36-43% of the xylem) predominate over vessel conduits (22-28%) and living parenchyma cells (26-28%). Conversely, in forbs, *Lychnis* and *Silene*, living parenchyma (both latewood tangential bends and multiseriate rays) predominate (50-56%) over lignified (15-20%) and conductive (24-26%) tissues. For grassland forbs, therefore, a combination of non-lignified latewood parenchyma bands and lignified fibers surrounding earlywood vessels assures both sugar storage capacity required for fast regeneration next spring as well as mechanical stability of aboveground shoots, respectively.

In addition to contrasting anatomical adaptations, differences in other functional traits (e.g., plant height, rooting depth) may contribute to explaining species-specific growth-climate relationships. Unlike *Thymus*, characterized by low-creeping stems and shallow roots (Kutschera & Lichtenegger, 1992), and susceptible to extreme summer heatwaves, the 30–60 cm tall and deep-rooted *Silene* was the only species whose growth was not affected by summer weather fluctuations but instead was contingent on late winter and early spring precipitation. Therefore, deep roots may prevent dislodging and promote drought resistance, potentially helping to overcome exceptional heatwaves and seasonally low water table (Schenk & Jackson, 2002). Also, Reader et al. (1993) found that the rooting depth in *Silene* seedlings increased with simulated soil dryness. We found a significant increase in radial growth in *Silene* with increased February and March precipitation, which probably creates sufficient soil water supplies that deeper roots can suck in during the summer drought to mitigate its harmful effect.

## 5 CONCLUSIONS

This study yielded several key findings and insights from exploring temporal changes in species-specific responses of plant population age structure and fitness to climate fluctuations. The focal species proved their ability to operate as biological data-loggers, yet with unique and distinct features. First, we found species-specific climate-recruitment relationships, probably due to differences in rooting abilities and seed mass. *Silene* and *Helianthemum* can use any wet period for seedling establishment, probably because of their deep roots; besides, they both produce larger seeds than the other two species, possibly pointing towards a whole-plant integrated regeneration syndrome. In contrast, *Thymus* and *Lychnis* recruitment appear less sensitive to climate fluctuation, probably due also to their very tiny seeds; their regeneration can be therefore more contingent on indirect effects, such as soil disturbance, gap formation, or mast seeding years.

Second, we found a strong correlation between growth and climate, indicating the ability of temperate dry grassland plants to react rapidly to changing climate. However, the observed relationships between growth and climate are species-specific and are probably related to differences in plant size, anatomy, and rooting depth. The growth of the shallow-rooted *Thymus* and *Lychnis* is sensitive to summer weather fluctuations, increasing with summer precipitation and decreasing with the summer heatwaves. In contrast, the deep-rooted *Silene* and *Helianthemum* are much less sensitive to summer weather, but both strongly depend on the late winter and early spring conditions. *Silene* is supported by warm and humid February and March, while *Helianthemum* favors drier and colder late winter and early spring conditions.

Third, we found senescing populations in all four species due to reduced recruitment in the warm and dry weather of the last decade. This may indicate a high vulnerability of the temperate dry grassland plant species to a combination of frequent summer heatwaves and prolonged droughts. Continuing trends of rising temperature and diminishing precipitation may result in species population contraction through impeded seedling recruitment. This can promote changes in community assembly, such as the spread of ruderal or non-native species (Gibson & Newman, 2019; Fischer et al., 2020). However, this can be partially offset by relatively high species longevity and slow growth, ensuring long-term population persistence.

The dependency of inter-annual recruitment and growth dynamics on multiple climatic drivers implies the crucial role of synergies and feedback mechanisms in the effects of temperature and precipitation. The different, species-specific responses of recruitment and growth to climate revealed here illustrate the challenges in drawing general conclusions about the effect of climate change without realizing how plants perceive these changes, i.e. a plants’ eye view (Liancourt et al., 2020). The differences in species strategies and responses to climate fluctuations belong to coexistence mechanisms that may ensure the long-term continuation of the grasslands. In periods of exacerbating climate change, some species may go extinct, but others will survive because of their different strategies.

## Supporting information

In-situ measured air temperatures and soil moistures

## ACKNOWLEDGEMENTS

J.D. and J.A. were supported by the Czech Science Foundation (Project: 21-26883S) and MSMT LTAUSA18007. G.O., L.C., F.E.M.C., and V.J. were supported by the Czech Science Foundation (Project: 19-14394Y). M.C. was supported by the Czech Science Foundation (Project: 19- 28491X). JK was supported by MSMT (LTT20003). J.D., J.A., J.K., G.O., L.C., F.E.M.C., and V.J. were supported by the Czech Academy of Sciences (RVO 67985939).

## AUTHOR CONTRIBUTION

J.D., J.A. and G.O. conceived and designed the study. G.O., L.C., and F.E.M.C. collected the data. V.J. analyzed root microsections. J.A. and J.D. analyzed the data. J.D., J.A., M.C. and G.O. led the writing of the paper. All authors contributed critically to draft revisions and gave final approval for publication of the paper. This research has not been previously presented elsewhere.

