## Supplementary material for "Climate fluctuations drive the recruitment and growth of temperate grassland plants": In-situ measured air temperatures and soil moistures

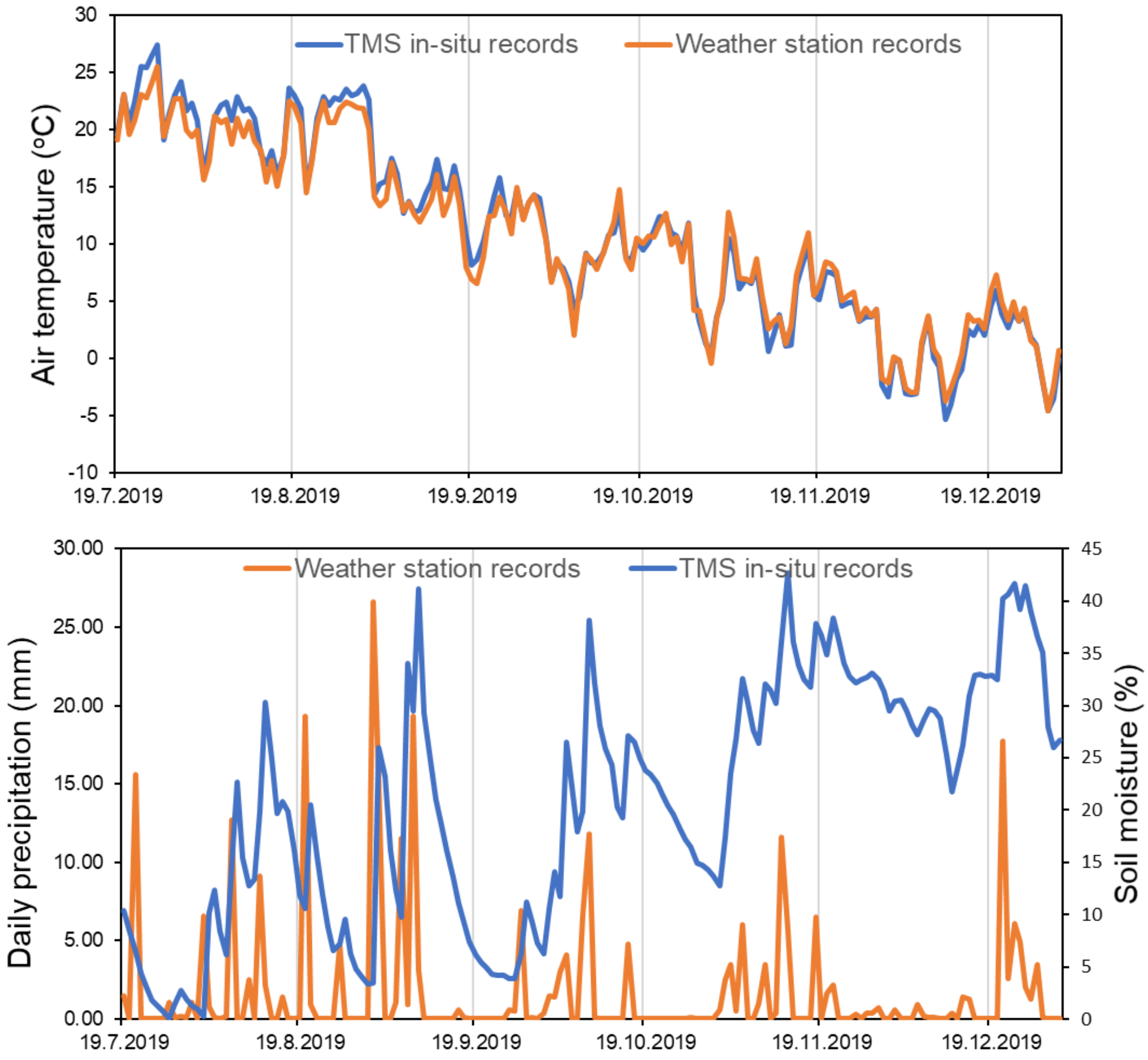


**Figure S1.** The course of the in-situ recorded mean daily air temperature at 10 cm aboveground and soil moisture at 2-10 cm belowground (using TMS loggers, blue curve), compared with the temperature and precipitation records from the nearby weather station in Velké Meziříčí (E 16.0086; N 49.3528, orange curve).
